# Striking Departures from Polygenic Architecture in the Tails of Complex Traits

**DOI:** 10.1101/2024.11.18.624155

**Authors:** Tade Souaiaia, Hei Man Wu, Anil P.S. Ori, Shing Wan Choi, Clive J. Hoggart, Paul F. O’Reilly

**Affiliations:** Department of Cellular Biology, Suny Downstate Health Sciences University, Brooklyn, NY, USA; Department of Genetics and Genomic Sciences, Icahn School of Medicine, Mount Sinai, NY, NY, USA; Department of Psychiatry, Amsterdam UMC, University of Amsterdam, Amsterdam, The Netherlands

**Author notes:** These authors contributed equally to this work (Tade Souaiaia, Hei Man Wu).

## Abstract

Understanding the genetic architecture of human traits is of key biological, medical and evolutionary importance[1]. Despite much progress, little is known about how genetic architecture varies across the trait continuum and, in particular, if it differs in the tails of complex traits, where disease often occurs. Here, applying a novel approach based on polygenic scores, we reveal striking departures from polygenic architecture across 148 quantitative trait tails, consistent with distinct concentrations of high-impact rare alleles in one or both tails of most of the traits. We demonstrate replication of these results across ancestries, cohorts, repeat measures, and using an orthogonal family-based approach[2]. Furthermore, trait tails with inferred enrichment of rare alleles are associated with more exome study hits, reduced fecundity, advanced paternal age, and lower predictive accuracy of polygenic scores. Finally, we find evidence of ongoing selection consistent with the observed departures in polygenicity and demonstrate, via simulation, that traits under stabilising selection are expected to have tails enriched for rare, large-effect alleles. Overall, our findings suggest that while common variants of small effect likely account for most of the heritability in complex traits[3], rare variants of large effect are often more important in the trait tails, particularly among individuals at highest risk of disease. Our study has implications for rare variant discovery, the utility of polygenic scores, the study of selection in humans, and for the relative importance of common and rare variants to complex traits and diseases.

## Introduction

Genetic architecture describes the number, effect sizes and allele frequencies of genetic variants affecting a complex trait^[1]^. Extensive efforts have been made to elucidate the genetic architecture of human traits over many decades and across multiple fields, including population genetics^[4]^, quantitative genetics^[5]^ and genetic epidemiology^[6]^. Recent genome-wide association and sequencing studies have identified many thousands of common and rare variants associated with a wide range of complex traits^[6–8]^. The number of variants identified is now so large that a complete picture of the genetic architecture of many complex traits is starting to emerge. Recent publications illustrate all known variants for a trait on a spectrum of allele frequency versus effect size, producing a characteristic “trumpet” shape^[9,10]^ that reflects the influence of natural selection in shaping the genetic architecture of human traits^[4,11]^. Stabilising selection, a key driver of genetic variation^[4,12]^, reduces fitness in individuals whose trait values deviate from an optimal mean, which consequently limits the frequency of large-effect alleles. Despite this, most statistical genetics methods suggest that common, small-effect variants jointly account for the majority of trait heritability^[3,13,14]^. However, little is known about how genetic architecture varies across the trait continuum. For example, it is unknown whether common and rare alleles are evenly spread through quantitative trait distributions, or if individuals in the trait tails (e.g. the lowest and highest 1% of LDL, BMI, glucose) are more likely to harbour high-impact, rare alleles.

We hypothesise that the strength and type of selection acting on a complex trait influences the distribution of common and rare alleles along the trait continuum. Specifically, we propose that many complex traits have experienced sufficiently strong selection to generate marked enrichments of large-effect, rare alleles in the trait tails (***Fig. 1***). This concept contrasts with the classical infinitesimal model, initially proposed by R.A. Fisher in 1918^[15]^, which posits that quantitative traits result from the cumulative effects of numerous alleles, each one of tiny effect. This “polygenic” model has since been extended to explain genetic liability of complex diseases and has proven incredibly effective in describing the genetic architecture of complex traits^[16,17]^. Indeed, many widely used statistical genetics methods rely on its assumptions^[18–20]^. Our hypothesis diverges from the infinitesimal model to the extent that selection constrains large-effect alleles to remain at low frequencies, effectively concentrating rare alleles in the tails of complex traits (***Fig. 1***). Thus, while the infinitesimal model accurately reflects the genetic architecture for the majority of the population, where polygenic burden predominates, it may be a poor approximation in trait tails subject to negative selection, where genetic architecture is more heterogeneous. This would be a key caveat to the infinitesimal model because in this case rare variants may make a modest contribution to overall trait variance^[3]^, but a substantial contribution in trait tails, which are of critical evolutionary and clinical significance^[21–23]^.

**Fig. 1:**
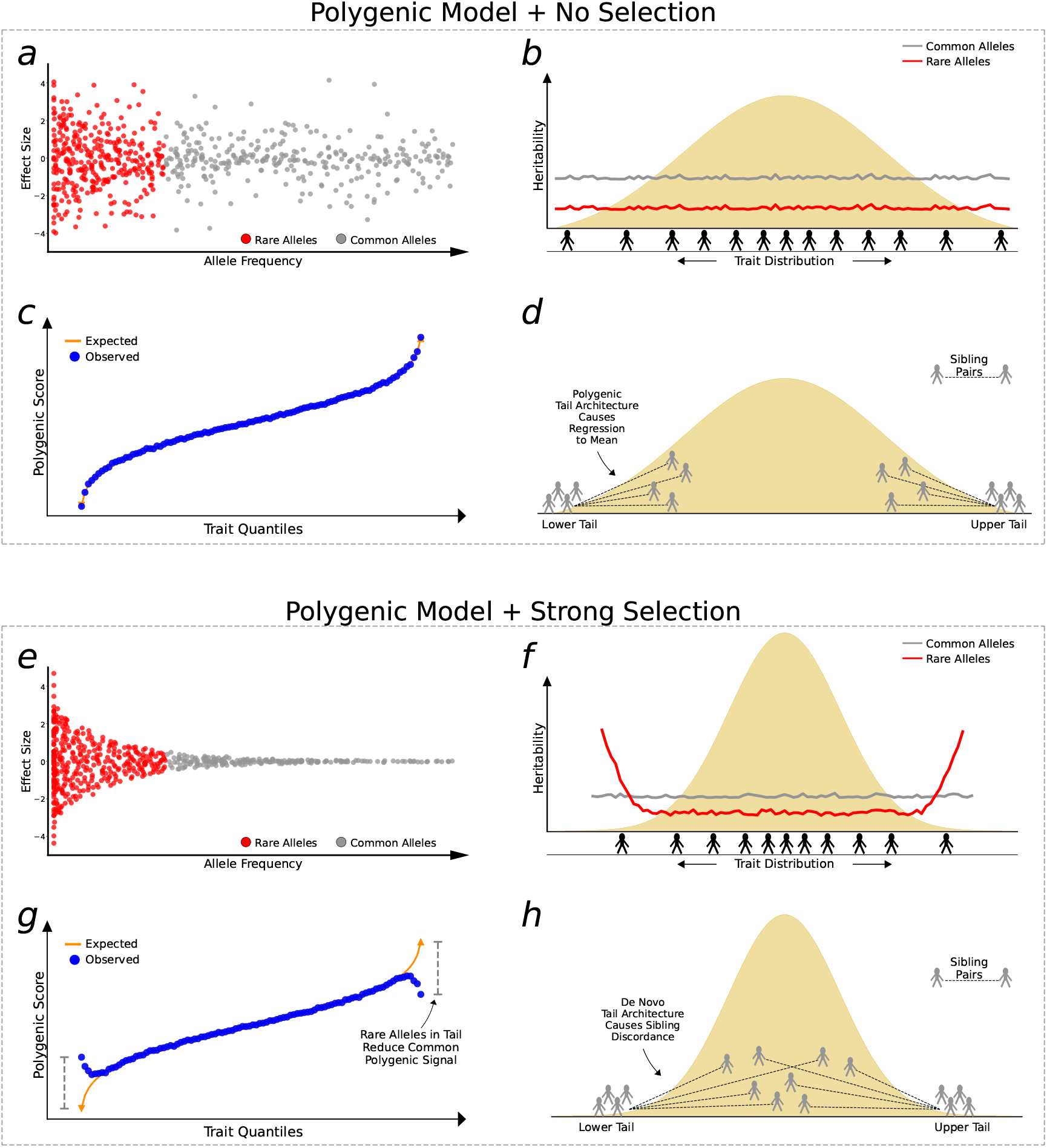
Two polygenic models of complex trait genetic architecture. In the polygenic model with no selection: **a**, the minor allele frequency (MAF) and effect size of variants affecting the trait are uncorrelated, **b**, common and rare variants are uniformly distributed across the trait continuum and common variants explain more heritability, **c**, the trait and polygenic score have a linear, monotonic relationship, and **d**, siblings of individuals in the tails of the trait distribution also show low/high trait values, with regression-to-the-mean dependent on trait heritability ^[2]^. The hallmark of this model is that polygenic architecture predominates the entire trait distribution. In the polygenic model with strong selection (here, stabilising selection): **e**, there is a strong relationship between MAF and effect size, whereby only rare alleles have large effect, **f**, common variants explain more heritability overall, but rare variants contribute more in the tails, **g**, the polygenic score (derived from common variants) regresses to the mean in the trait tails due to enrichment of high-impact rare alleles not included in the polygenic score, **h**, assuming selection is so strong to produce tails characterised by *de novo* architecture, then siblings of individuals in the trait tails have trait values drawn from the background distribution. The hallmark of this model is that polygenic architecture prevails across most of the trait distribution, except for the tails where rare-variant architecture predominates. These models illustrate two extremes of selection, neither of which will be applicable for most traits (indeed recent work suggests polygenicity may be caused by selection ^[4]^); however, most complex traits should be captured by an intermediate model that lies somewhere between these two extremes.

The existence of pathogenic alleles with dramatic effects on quantitative traits is already known in isolated cases. For example, mutations in ACAN result in short stature^[24]^, mutations in LEP cause leptin deficiency and severe early onset obesity^[25]^, while those in HNF1A produce large deviations in glucose homeostasis and early-onset diabetes^[26]^. Therefore, it is already known that there are single alleles of sufficient effect size to generate extreme trait values in individuals. Moreover, Fiziev et al.^[27]^ demonstrated that rare polygenic risk scores, calculated based on the PrimateAI-3D algorithm, were more predictive for individuals with extreme, rather than average, trait values. However, so far there has been no systematic investigation into how genetic architecture varies across complex trait distributions and whether, in particular, the tails of quantitative traits have distinct genetic architecture.

In this study, we apply multiple approaches to interrogate how genetic architecture varies across the trait continuum. First, we introduce the PRS-On-Phenotype outlier (“POPout”) test, a novel method that uses polygenic risk scores^[20]^ (PRS) to detect deviations in polygenic expectations in the tails of trait distributions. The premise of the POPout test is that a PRS derived from common variants should exhibit a positive, linear correlation with its corresponding trait (***Fig. 1c***). However, if rare alleles are concentrated in the trait tails, then the PRS is expected to regress towards the mean in these extremes (***Fig. 1g***). We first apply POPout to 74 quality-controlled quantitative traits in individuals of European ancestry from the UK Biobank^[28]^, followed by replication in individuals of multiple ancestries from the UK Biobank and in the US-based All of Us cohort^[29]^. Next, we employ an independent method^[2]^ that uses sibling trait variability to infer the genetic architecture of trait tails (***Fig. 1d, Fig. 1h***) in over 17k sibling pairs from the UK Biobank. This sibling-based approach provides complementary evidence to POPout, since it leverages non-genetic family data to infer tail architecture. Across these approaches and cohorts, we observe widespread and substantial departures from polygenic expectations in trait tails (***Fig. 2, Fig. 3***). To gain insight into the causes of these observations, we investigate whether natural selection may drive these findings (***Fig. 4***). Extending a method^[30]^ that uses lifetime reproductive success to infer ongoing positive, negative, and stabilising selection, we find consistency between the type of selection inferred and the tail(s) of a trait that appear to be selected against according to our POPout results. Supporting this, simulations using the forward-in-time simulator SLiM^[31]^, show that stabilising selection results in POPout effects in both tails, whereas traits simulated under neutrality display no such effects. Finally, we observe that trait tails with POPout effects are associated with a higher frequency of significant exome-based rare variants, reduced fecundity in both males and females, and advanced paternal age. Our findings reveal the distinct nature of genetic architecture in the tails of complex traits, highlighting the importance of distinguishing between the diverse genetic profiles that can lead to extreme trait values in different individuals and in different traits.

**Fig. 2:**
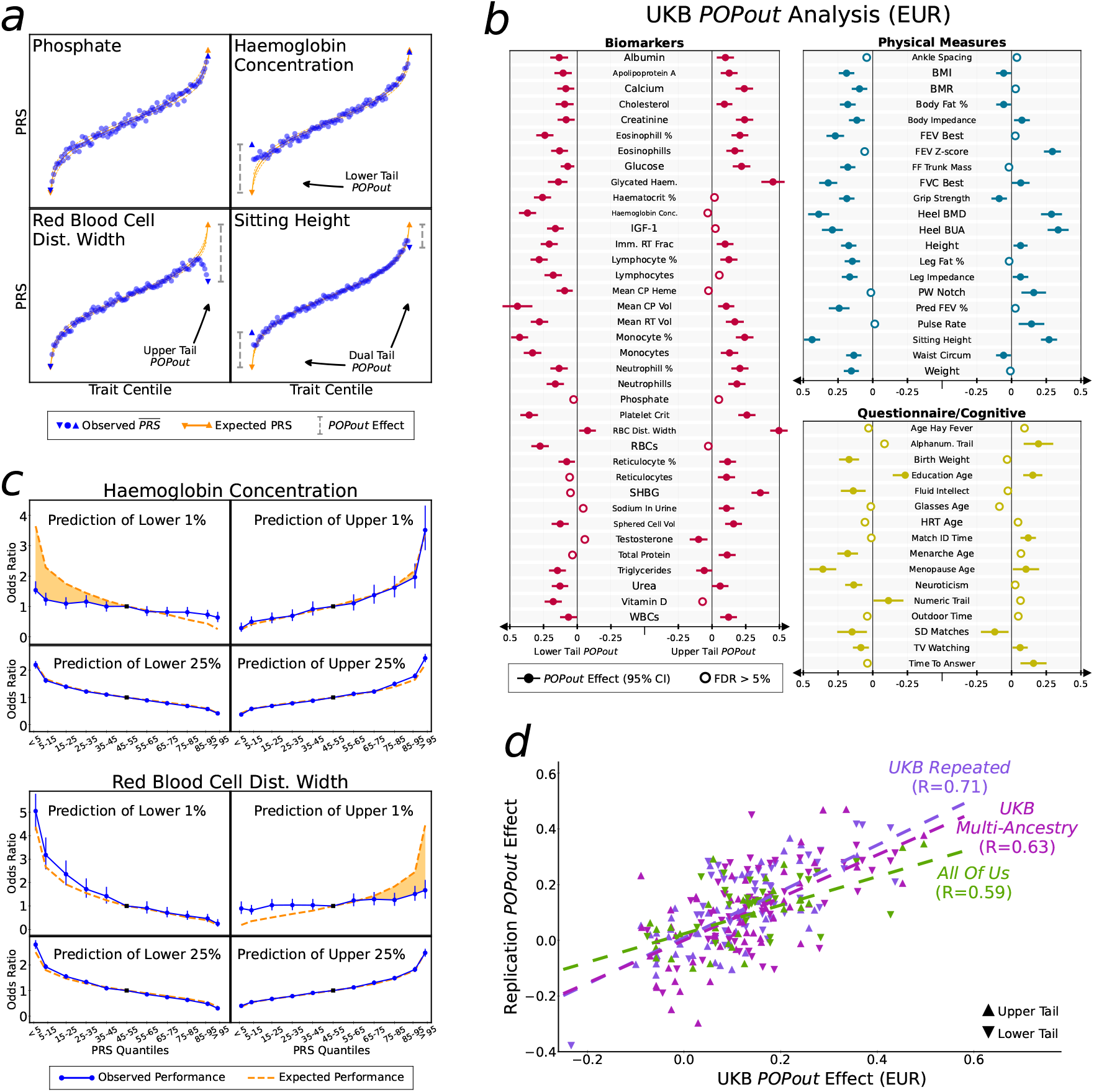
Population-based POPout analyses. **a**, Trait quantiles (1% increments) plotted against mean PRS value for four traits with different POPout characteristics. Expected PRS (orange) calculated from regression of PRS on trait. **b**, POPout results across 74 QC’ed traits in UKB EUR data. Effects with 95% confidence intervals shown. **c**, Odds ratio corresponding to the lowest/highest 1%/25% of the trait as index PRS quantile versus the central PRS quantile. Orange dashed line is expected performance assuming infinitesimal model (shaded area highlights the difference with observed). **d**, POPout effect sizes from UKB EUR analysis (of b) plotted against replication POPout effect sizes for upper and lower tails of all overlapping traits. Correlation between UKB EUR and replication effect sizes calculated using Pearson.

**Fig. 3:**
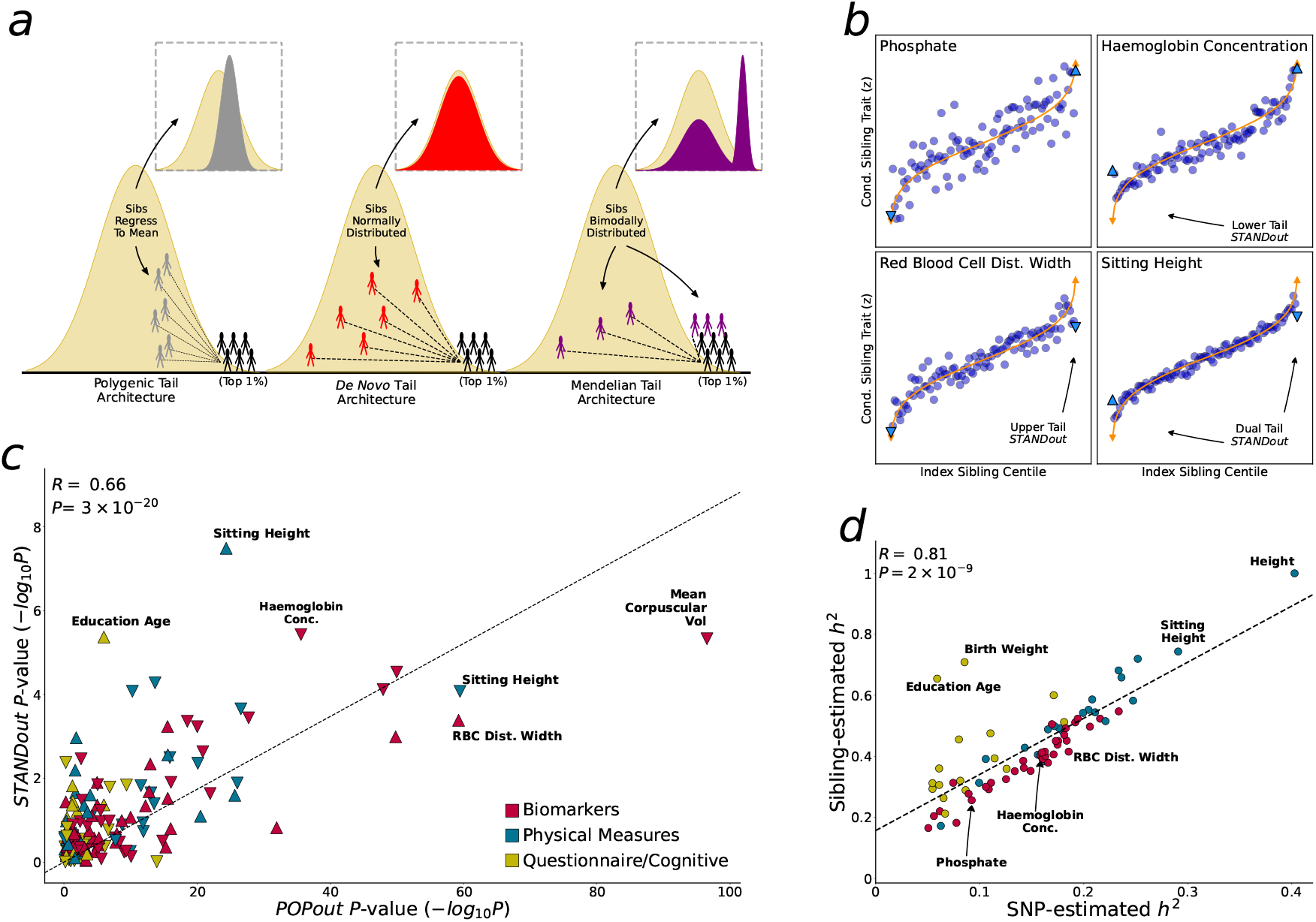
Tail architecture inferred from siblings. **a**, Schematic of relationships between sibling trait values under tails governed by polygenic (grey), *de novo* (red) and Mendelian (purple) architecture, where Mendelian here refers to rare (but not *de novo*) high-impact alleles segregating in families. **b**, Index sibling trait quantile (1% increments) plotted against mean trait value of other (conditional) sibling for the four traits selected with different POPout characteristics (see Fig.2a). **c**, POPout *P* -values plotted against STANDout *P* -values for the 74 traits. Correlation and significance calculated using Pearson. **d**, LD score regression ^[19]^ SNP-estimated heritability plotted against heritability estimated from sibling pair trait similarity ^[2]^.

**Fig. 4:**
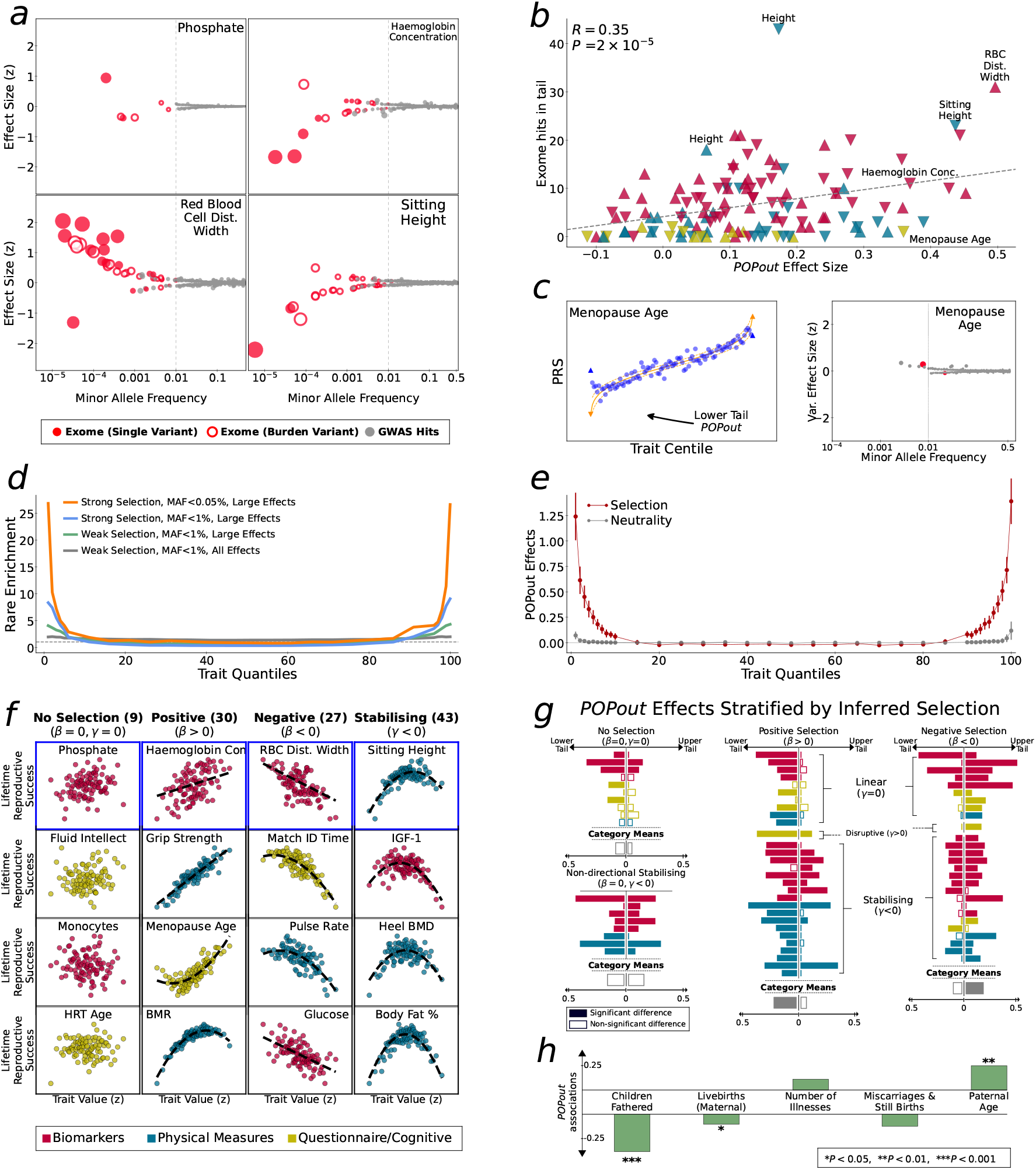
Rare variants and natural selection as putative causal mechanisms of observed POPout effects. **a**, Trumpet plots ^[10]^ (minor allele frequency vs effect size) of significant variants reported in major exome ^[34]^ and GWAS ^[35]^ studies for the four traits selected for their different POPout effects (see Fig.2a). **b**, POPout effect size vs number of significant exome variants ^[34]^. For upper tail POPout effects, the number of trait-increasing rare alleles are counted, while for the lower tail the number of trait-reducing rare alleles are counted. Correlation and significance calculated using Pearson correlation. **c**, PRS-on-trait quantile plot (left) and trumpet plot (right) for self-reported age of menopause in UKB. **d**, Enrichment of rare vs common alleles in simulations under weak and strong selection scenarios, stratified by rare variant classification and effect size (see Methods). **e**, POPout effects across trait quantiles averaged over 100 simulations (mean and CIs shown) under neutrality and selection. **f**, Trait value vs relative lifetime reproductive success for 16 example traits. Best fitting model (dashed line) determined by BIC (*β* = linear, *γ* = quadratic, regression parameters) used to classify selective pressure acting on trait. **g**, POPout effect sizes for traits are grouped by selection category inferred by BIC model (panel **f**, models with linear and quadratic terms are categorised by direction of linear term). Differences between lower and upper tail effects tested by two-sample *t*-test in each of the four categories). **h**, Association between POPout effect size and five traits relating to fitness. For each proxy fitness trait, Spearman rank correlation was calculated between the mean of the trait in individuals in the upper and lower 1% trait tails and POPout effect size for the corresponding tail across the 74 traits.

## Results

### Tail architecture inferred by PRS

Here we first investigate how genetic architecture varies across the trait continuum in a range of quantitative traits using UK Biobank (UKB) genotype-phenotype data^[28]^. We consider traits from the broad UKB trait categories of *Biomarkers, Physical Measures, Touchscreen Questionnaire*, and *Cognitive Function*. After quality control (QC) steps (see Methods), to ensure, for example, that the traits selected for analysis are heritable, approximately normally distributed and sufficiently independent, 74 traits remain for analysis. To limit the contribution of environmental factors, we residualised the traits with a range of covariates, including age, sex, smoking, alcohol intake, menopause status; we also removed individuals with cancer diagnoses or who were pregnant, as well as individuals on statins or on insulin medication within the biomarker category. After standard QC of the genotype data there were 369,132 individuals of European (EUR) ancestry available for the primary analysis.

We have developed a method that leverages polygenic risk scores (PRS), derived from common variants, to infer genetic architecture in the tails of quantitative traits. In standard PRS analyses^[20]^, PRS is a predictor of outcome, with results often displayed as a quantile plot showing higher values (or odds ratio) of the outcome with increasing PRS quantiles^[32,33]^. Here we invert this relationship, so that trait is a predictor of PRS, displaying results as trait quantiles on the X-axis and PRS on the Y-axis. This simple reversal of the usual relationship enables us to investigate how the PRS varies along the trait continuum. Under a null hypothesis corresponding to a polygenic model^[17]^, with environmental factors adjusted or jointly normally distributed, we expect a linear relationship between the trait and PRS (***Fig. 1c***). In contrast, if the tails are enriched for rare alleles of large effect, not included in the PRS derived from GWAS on common variants, then the average PRS in the lower and upper quantiles will regress to the population mean PRS (***Fig. 1g***). We formally tested for deviations from polygenic expectations by performing *t*-tests on the residuals of a regression of PRS on trait. Each of the middle 80 trait centiles were tested as a QC step here, while the analysis test - which we refer to as the “PRS-On-Phenotype outlier test”, or POPout test - is applied to the lower and upper trait centiles (see Methods). The POPout test is applied to the bottom and top 1% of trait values here since these quantiles are considered sufficiently extreme to be subject to selection, given the typical prevalence of complex disease, yet broad enough for sufficient sample size. We applied the POPout test to the 74 traits in the UK Biobank, splitting the sample of 369,132 individuals into two halves, in which a GWAS is performed in the first half and used to derive PRS in the second half for the analysis. Figure 2a displays PRS-on-trait quantile plots corresponding to four of the traits that illustrate different types of observed relationships. Phosphate shows a relationship that follows polygenic expectations (orange line) closely. Haemoglobin concentration shows a strong departure from polygenic expectations in the lower tail, corresponding to a POPout effect size of 0.37 (*P* = 2.6 × 10^−36^), consistent with strong enrichment of rare alleles in the lower tail, while red blood cell distribution width shows a strong regression-to-mean of PRS in the upper tail (POPout effect: 0.5; *P* = 5.7 × 10^−60^), consistent with a concentration of rare alleles of large effect residing in individuals of the upper tail. Sitting height shows strong POPout effects in both tails (lower tail *P* = 3.5 × 10^−60^, upper tail *P* = 4.3 × 10^−25^), consistent with rare allele enrichments in both tails and the action of strong stabilising selection. The four traits included in Figure 2a were selected because they provide clear illustrations of the different types of relationship and are not reflective of the general results. In the full POPout analysis of 74 traits, 68 traits had significant (FDR*<* 5%) POPout effects in one or both tails, suggesting pervasive departures from polygenic architecture in the tails of complex traits in general (***Fig. 2b, Supplementary Table 1***). Of the 107 tails with significant POPout overall, 97 showed POPout effects in the direction expected according to rare allele enrichment in the tails, offering support to our hypothesis, while numerous trait tails showed striking departures from polygenicity. The 10 significant negative POPout effects could be a result of epistatic effects or large, non-Gaussian, unadjusted environmental risk factors (see Discussion).

To exemplify the impact of these observed deviations from polygenic expectations, we emulated PRS prediction of case/control status for diseases hypothetically diagnosed in individuals with low haemoglobin and high red blood cell distribution width, with prevalences of 1% and 25% (***Fig. 2c***). While prediction of case/control status using the trait PRS is as expected (orange dashed line) for a disease prevalence of 25%, the PRSs underperform substantially for both traits for disease prevalence 1%. For haemoglobin, the mean odds ratio (OR) for individuals in the bottom 5% of PRS is expected to be 3.7 but is in fact 1.8, while, for red blood cell distribution width, the mean OR for individuals with top 5% PRS values is 1.8 instead of an expected 4.1.

### Replicating POPout Effects

Here we performed three different forms of replication to test the generalizability of our primary findings that used UKB European ancestry data. In each replication dataset, we only included traits that passed our POPout QC (see Methods) and have sample size over 5k individuals based on inferred power (see ***Supplementary Fig. 1***). First, to investigate if POPout effects of the primary analysis are due to residual non-genetic factors not captured by our QC or covariate adjustments, we repeated the analysis in a held-out sample of individuals that had traits measured at baseline and at approximately 5-year follow-up. By taking mean values of the two repeat measures, the impact of transient, environmentally influenced factors, such as infections, short-term medication use, or diet, should be minimised. This ‘UKB Repeated’ analysis, performed on up to ∼ 15k individuals across the 50 (of 63 overlapping) traits that passed QC, showed a strong and significant (*R* = 0.71, *P* = 2.3 × 10^−16^) correlation with the results of the primary analysis (***Fig. 2d***), suggesting genetic contribution to the observed POPout effects of the primary analysis. Next, we investigated the generalizability of our findings to a held-out sample of UKB multi-ancestry ancestry individuals (see Methods). This ‘UKB Multi-Ancestry’ analysis, performed on up to ∼ 22k individuals across the 54 traits that passed QC, produced highly correlated and significant replication (*R* = 0.63, *P* = 3.7 × 10^−13^) of the primary results (***Fig. 1d***). Finally, we apply the same approach to European ancestry individuals from the US-based All of Us data^[29]^ to evaluate the generalizability to a matched-ancestry sample but from a clinical-based cohort from a different continent, with different collection and assaying procedures, and a different healthcare system and set of environment factors. This ‘All of Us’ analysis, performed on up to ∼ 127k individuals across the 28 (of 33 overlapping) traits that passed QC, also showed strong replication (***Fig. 1d***) of the primary analysis results (*R* = 0.59, *P* = 1.6 × 10^−6^). See Supplementary Material (***Supplementary Fig. 1***) for a full breakdown of all replication analyses.

### Tail architecture inferred from siblings

Next, we seek to validate our findings further using family data. We have developed a method^[2]^ that evaluates trait data in large samples of siblings to infer genetic architecture in the tails of quantitative traits. This provides an alternative, complementary approach to infer tail architecture from independent, family-based, non-genetic data. The intuition of the approach (***Fig. 3a***) is that individuals with extreme trait values have siblings whose trait values reflect the architecture in the tails, such that under: (1) polygenic architecture: siblings have high but less extreme trait values, (2) *de novo* architecture: siblings have trait values reflective of the general population, (3) Mendelian architecture (i.e. due to rare, high-impact alleles that segregate in families): siblings either have similarly extreme trait values or have trait values that reflect the general population.

Having established theoretical expectations of sibling similarity according to genetic architecture, we developed tests to distinguish between these three possibilities, which we extend here to a joint test, which we refer to as STANDout (Methods). STANDout infers general departures from polygenic expectations in trait tails from trait-only data and is powered to identify tail architecture enriched for large-effect, rare alleles (that can cause individuals to ‘stand out’ from their siblings for given traits), assuming environmental factors are adjusted or jointly normally distributed.

The UK Biobank (UKB) includes over 17k sibling pairs of European ancestry available for our STANDout analyses, none of which were included in our population-based analyses. There were too few sibling pairs from the multi-ancestry sample for a sufficiently powered multi-ancestry sibling analysis. Data across all 74 traits were available for analysis. Replicating our POPout analysis, our STANDout analysis showed no departure from polygenic expectations for phosphate, showed a significant lower tail STANDout effect for haemoglobin concentration (*P*_*lower*_ = 2 × 10^−5^), a significant upper tail STANDout effect for red blood cell distribution width (*P*_*upper*_ = 4 × 10^−4^), and significant effects in both tails for sitting height (*P*_*lower*_ = 7 × 10^−4^, *P*_*upper*_ = 3 × 10^−8^; see ***Fig. 3b*** and ***Fig. 2a*** for comparison). More broadly, our sibling-based STANDout analyses replicated our population-based POPout analyses across the 74 traits well (***Fig. 3c***; *R* = 0.66, *P* = 3 × 10^−20^). This is reassuring given that the two methods use such orthogonal approaches and types of data. Further reassurance of the validity of our sibling-based method is provided by the strong correlation (*R* = 0.81, *P* = 2 × 10^−9^) between SNP-based heritability (*h*^2^), estimated by the popular LD score regression approach^[19]^, and the sibling-based *h*^2^ estimated as a key parameter in the STANDout approach (***Fig. 3d***).

### Selection shapes tail architecture

In previous sections, we observed and replicated widespread departures from polygenic expectations in the tails of quantitative traits. Here, we investigate evidence for the role of rare variants and selection in causing these patterns (***Fig. 1***). Backman et al.^[34]^ conducted the most comprehensive trait-wide analysis of UK Biobank exome sequence data to date, and here we utilize their results to investigate whether traits with significant tail departures are associated with a greater number of significant exome variants (or ‘hits’), with effects in the expected direction. Considering our four example traits, phosphate has two exome hits with positive and four with negative effect sizes, haemoglobin concentration has eight with positive and twelve with negative effect sizes, red blood cell distribution width has twenty-eight with positive and five with negative effects, and sitting height has ten with positive and twenty with negative effects (***Fig. 4a***). Across the 74 traits, we found that POPout effect sizes from our primary analysis are significantly positively correlated (*R* = 0.35, *P* = 2 × 10^−5^) with the number of exome hits with effects in the direction corresponding to the tail of the POPout effect (***Fig. 4b***). This provides direct support for the observed complex tail architecture being underlain by rare alleles. Findings by Fiziev and colleagues^[27]^ offer further support. The authors derived rare PRS across a range of quantitative traits in the UKB and found that they performed particularly well in predicting individuals with extreme trait values. However, we highlight one result that suggests that rare PRS may perform poorly for traits under extremely strong selection. The trait with the most significant POPout effect in the questionnaire/cognitive category (***Fig. 2b***) was *Age at Menopause*, which had an effect consistent with rare allele enrichment in the lower tail; yet the exome data showed no hits of negative effect (***Fig. 4c***). However, given the rationale for early age of menopause being under particularly strong negative selection, it may be that only ultra-rare alleles of large effect remain in the population, producing a PRS regression-to-the-mean in the tail but little signal in exome data.

To gain greater insight into how selection may impact variation in genetic architecture across the trait continuum, we performed simulations using the forward-in-time population genetic simulator SLiM^[31]^. Using standard population genetic parameter settings (Methods), we simulated genetic variation data and a polygenic quantitative ‘trait’ calculated as the sum of genome-wide alleles, with effect sizes drawn from a Gamma distribution. After an initial burn-in period under neutrality, the trait was subjected to stabilising selection or strong stabilising selection. After stabilising selection, rare alleles were enriched in the trait tails compared to under neutrality, and this effect was more pronounced for very rare (MAF < 0.05%) alleles and after strong stabilising selection (***Fig. 4d***). Under neutrality, approximately 1% of simulated individuals in the top trait centiles harbored rare alleles of sufficiently large effect to cause their extreme trait value, whereas under strong stabilising selection about 28% of individuals in the tails carried such alleles. As a consequence, the trait tails showed a marked difference between trait value and common PRS as averaged across 100 simulation replicates (***Fig. 4e***), mimicking the POPout effects of the real data analyses. These simulations suggest that selection can have a critical impact on how genetic architecture varies across the trait continuum and can generate substantial rare-variant architecture in the tails of traits that are otherwise governed by polygenic burden.

To test this more explicitly in our real data, we applied a method^[30]^ for inferring ongoing selection from “lifetime reproductive success” to our 74 traits in the UKB data (see Methods). We use Bayesian Information Criteria (BIC) to distinguish “no selection”, “positive”, “negative”, and “stabilising” selection in the fitted models (***Fig. 4f***). Next, we stratified the POPout results of the primary analysis into groups of traits according to the categories of inferred selection and tested if the POPout effects reflect the inferred selection (***Fig. 4g***). Despite the snapshot into historical selection provided by contemporary reproductive success, POPout effects were reflective of inferred selection. There were no significant differences in mean POPout effect between lower and upper tails that were inferred to be under no selection (lower mean = 0.09, upper mean, 0.04, *P* = 0.18) and non-directional stabilising selection (lower mean = 0.18, upper mean = 0.14, *P* = 0.52). As expected there were significantly larger POPout effects in the lower tail for traits with inferred positive selection (lower mean=0.2, upper mean=0.04, *P* = 7 × 10^−7^) and significantly larger POPout effects in the upper tail for traits with inferred negative selection (lower mean = 0.08, upper mean = 0.17, *P* = 0.02), Finally, we tested for associations between POPout effects and outcomes that reflect proxy measures of fitness (***Fig. 4h***), finding that POPout effects are significantly negatively correlated with number of offspring (paternal: *P* = 0.0001; maternal: *P* = 0.0233), significantly positively correlated with paternal age (*P* = 0.0096) and not significantly associated with self-identified illnesses or number of miscarriages and still births.

## Discussion

Our study is the first dedicated investigation into how genetic architecture varies across human complex trait distributions. Applying our novel PRS-based method, POPout, and family-based method^[2]^, STAND-out, to 74 quantitative traits of the UK Biobank, we observed widespread departures from polygenic expectations in one or both tails of most of the traits. Our replication results suggest that these deviations from the infinitesimal model in complex trait tails are a stable, replicable, global phenomenon. Interrogating the causes of these observations, we found direct evidence for the involvement of high-impact, rare variants from large-scale exome data^[34]^, adding to evidence from recent research demonstrating the strong performance of rare PRS in prediction of extreme trait values^[27]^. Finally, several lines of investigation pointed to selection as a key mechanism driving the observed signals. Firstly, forward-in-time population genetic simulations^[31]^ showed enrichment of rare alleles of large effect in trait tails after a period of stabilising selection, as well as associated tail deviations in PRS that mimic real data POPout effects. Secondly, the type of ongoing selection that traits are subject to, as inferred by reproductive success^[30]^, was consistent with observed POPout effects; for example, traits inferred as subject to positive selection had significantly greater POPout effects in the lower tail than the upper tail. Thirdly, trait tails with POPout effects were associated with reduced overall fecundity in females and males, as well as advanced paternal age, which is a known source of increased *de novo* mutations^[36]^. Overall, these findings provide robust support for our overarching hypothesis: namely, that selective pressures on traits have led to pervasive concentrations of high-impact, rare alleles in the tails of many human traits, resulting in heterogeneous architecture that deviates markedly from the infinitesimal model^[17]^ in the tails.

Our work has several immediate practical and theoretical implications. Firstly, in order to optimise the discovery of rare variants in future studies on complex traits, researchers should first infer how genetic architecture varies across the trait distribution. Sequencing individuals with average PRS in the lower tail of haemoglobin concentration, for instance, is likely to yield greater discovery of rare variants than a design blind to POPout effects (***Fig. 2***a). Likewise, *a priori* evidence of enrichment of *de novo* mutations derived from our sibling approach^[2]^ would motivate trio sequence studies, rather than population sequence or genotype studies. Moreover, our results could be used to boost the power of gene-burden testing^[37]^ by assigning weights to individuals that reflect the prior probability of harbouring high-impact, rare alleles.

User-friendly software to implement our POPout and STANDout tests is openly available for such use and applicable to widely available population genotype and family trait data, respectively. Secondly, any study performing PRS analyses in relation to a quantitative trait, or disease with an informative biomarker, should consider conducting POPout analyses to aid interpretation of results, including PRS predictive accuracy, and facilitate improved prediction in the sample under study. Thirdly, we introduce and validate a new theoretical perspective on how selection contributes to shaping genetic variation. Recent appreciation of the extent of polygenicity across human traits has motivated a shift in focus from selective pressures acting at the variant or locus level^[38,39]^, to polygenic adaptation acting on the trait^[40,41]^. This has led to important advances in our understanding of how selection may have generated^[4]^, and can be inferred from^[42]^, widespread polygenicity. However, until now there has been no consideration of how different selective pressures on a trait can lead to different degrees of polygenicity across the trait continuum. Our novel insights into the impact of selection on genetic architecture should contribute to understanding of population genetic processes and can motivate the development of new theory to infer and evaluate historical selection on complex traits. Finally, the reduced predictive accuracy of PRS in trait tails with POPout effects indicates theoretical limits on future predictive performance of common PRS based for the relevant traits. Incorporating rare variants into PRS will improve prediction, but if much of the missing trait variance is explained by unidentified ultra-rare variants, then prediction will still be limited, particularly for individuals at greatest risk of deleterious outcomes. However, iterative application of our POPout approach can help to overcome this: POPout results can be used to increase rare variant discovery, as described above, and then newly discovered rare variants can be integrated into PRS, increasing PRS accuracy and, in turn, the power of subsequent POPout analyses to identify residual unidentified rare variants.

Our study had a number of limitations and areas for future follow-up. Firstly, and most important, we cannot rule out the contribution of environmental factors in generating the observations in relation to complex tail architecture. In fact, despite our focus on genetics here, we expect the environment to have made significant contributions to the observed departures from our null expectations. However, given control of major environmental risk factors, replication across multiple measures and cohorts, association with reported exome variants, and evidence for selection generating such signals from real and simulated data, enrichment of high-impact rare alleles is likely a major cause of our findings. Future work should interrogate potential genetic and environmental causes of extreme trait values at the individual level, with greater trait-specific attention than possible from our trait-wide analyses. Secondly, in our multi-ancestry replication analysis we combined individuals of many diverse ancestries into a single group, which poorly reflects global diversity and prevents evaluation of potential heterogeneity across populations. However, relatively large sample sizes are required to study tails of trait distributions in this way and so we chose to perform this grouping to establish whether our findings were valid across ancestries. Future studies should expand across global ancestries. Thirdly, we did not explicitly integrate rare PRS to recover observed reductions in absolute PRS in the tails. This was due to the recent publication of a highly comprehensive study that performs a similar analysis and provides direct support for our findings^[27]^, as well as the challenge in verifying observed effects that are due to presently unidentified ultra-rare and *de novo* effect alleles. Fourthly, we only utilised siblings given data availability, but our sibling-based method can be extended to other family members^[2]^ and applied accordingly. Fifthly, we did not consider pleiotropy between traits, which undoubtedly has a key role in selective pressures on individual traits^[42]^. Considering pleiotropy in the light of our findings to, for instance, determine whether there are stronger pleiotropic effects among individuals with inferred high-impact rare alleles versus those without, will be an extremely interesting area for follow-up; however, our conclusions should be valid irrespective of pleiotropy. Sixthly, we did not consider disease explicitly here due to the reliance of our POPout method on application to a continuous outcome. However, an informative biomarker or known risk factor of a disease can be used instead, and, critically, this provides a novel strategy for discovery of rare alleles with large effects on disease. Finally, selection was inferred from real data using data on fecundity, which only offers a crude proxy of selective pressures on a trait and is vulnerable to reverse causation. Despite this limitation, we observed significant association between selection inferred from fecundity and POPout effects, although future work should leverage the numerous methods for inferring selection from population genetic data for further verification.

This study devised and applied multiple approaches to interrogate how genetic architecture varies across the trait continuum, revealing striking and widespread departures from polygenic expectations in the tails of complex traits. Our findings offer a new perspective on genetic architecture and on the role of selection in shaping how contemporary genetic variation underpins human phenotypic variation. Future work can build on this to boost the discovery of rare variants, increase the accuracy of polygenic scores, infer historical selection from variation in genetic architecture across the trait continuum, and explicitly infer the relative contributions of different genetic and environmental causes of extreme trait values and disease in individuals.

## Acknowledgements

We thank the participants of the UK Biobank and the All of Us Research Program, as well as the scientists involved in the construction of these resources. This research has been conducted using the UK Biobank Resource under application 18177 (Dr O’Reilly). All participants gave full informed consent. This work was supported by NIH grants R01MH122866 and R01HG012773 and through the computational resources and staff expertise provided by the Data Ark and Scientific Computing teams at the Icahn School of Medicine at Mount Sinai. We are grateful for helpful discussions about this work with: Alanna Cote, Carla Giner-Delgado, Conrad Iyegbe, Judit García-González, Lathan Liou, Laura Sloofman, Louise Wain and Michael Weedon.

## Methods

### Data Processing

#### UK Biobank Genetic Data

The UK Biobank (UKB) is a prospective cohort study of approximately 500,000 participants recruited across the United Kingdom from 2006 to 2010^[43]^. Phenotype data of anthropometric, biological and lifestyle measures were collected at baseline and in follow-up surveys, with further linkage to health and disease record data. The genetic dataset consists of 488,377 samples genotyped at 805,426 SNPs. To define population ancestries, 4-means clustering analysis was conducted on the first two principal components (PCs) of the genotype data. Samples were then projected onto 1000 Genomes PCs to infer ancestry, resulting in 461,931 European, 11,074 South Asian, 7,935 African, 2,585 West Asian, 2,550 East Asian, and 1,619 individuals of other ancestries.

Subsequent to clustering, standard quality control (QC) procedures were applied independently to each ancestry cluster. SNPs with a minor allele frequency (MAF) *<* 0.01, genotype missingness > 0.02, or Hardy-Weinberg equilibrium test *P* -value *<* 10^−8^ were excluded. Samples exhibiting high levels of missingness or heterozygosity, inconsistencies in genetic-inferred and self-reported sex, or displaying aneuploidy of the sex chromosomes, were removed in accordance with recommendations from the UKB data processing team^[44]^. For the analyses in the general population, a greedy algorithm was employed to remove related individuals that maximises sample retention while removing all third-degree relatives (kinship coefficient > 0.044)^[45]^. After these QC steps, 411,948 unrelated individuals remained, of which 387,472 were of European ancestry. Included among the European ancestry cluster were 18,340 individuals with repeated measures that were set aside, along with 24,476 individuals of multiple non-European ancestries (see above), for replication analyses (below). This resulted in up to 369,132 unrelated European ancestry individuals for the primary POPout analyses (see below).

For the sibling analyses (see below), sibling pairs were selected by first identifying first-degree relatives, defined as those with kinship coefficient within the range 0.177 and 0.354 (expected value for siblings is 0.25) to account for the expected variation among siblings^[46]^. Only individuals of European ancestry were included in the sibling analyses due to insufficient statistical power of the multiple ancestries sample^[2]^. The proportion of SNPs with zero identity-by-state (IBS0) was used to distinguish sibling pairs from parent-offspring pairs, since parent–offspring pairs have IBS0≈0 across the autosomes. It has been shown in the UKB that there is good separation between pairs of first degree relatives at IBS0=0.00112^[46]^ and, thus, sibling pairs were selected as those exceeding this threshold. This provided a total of 17,289 sibling pairs for analysis.

### Quantitative Trait QC

#### Trait Selection

We considered all traits from the categories *Biomarkers* (453 traits), *Physical Measures* (271 traits), the *Touchscreen Questionnaire* (396 traits) and *Cognitive Function* (134 traits) to provide a broad group of well-studied, heritable traits. To minimise non-genetic causes of trait values, we then removed all individuals with a cancer diagnosis (34,698), on insulin treatment (1,311) or who were pregnant at baseline (109). We also removed 64,043 individuals on statins and other cholesterol-lowering drugs from the *Blood count* and *Blood Biochemistry* subcategories and 29,395 individuals on heart rate medication from the *Arterial stiffness* and *Blood pressure* subcategories. Next, to ensure that extreme outliers do not impact results, we removed all samples that had trait values six standard deviations or more from the mean. After these removals, we defined quantitative traits as those containing at least 10 distinct values and whose top two modal values comprise less than half the samples. Finally, we removed all traits with < 100, 000 samples remaining in our primary dataset of unrelated European ancestry individuals, resulting in a set of 180 quantitative traits. These traits were then residualised and subject to further QC as described below.

#### Trait Residualisation

To minimise the impact of environmental risk factors, trait values were residualised within each ancestry group (European and Multi-Ancestry) by applying a linear regression model to each trait, with covariates for *age, age*^2^, *age*^3^, sex, menopause status (among females), type-II diabetes status, coronary artery disease status, alcohol intake (Field ID: 1558), smoking (categorical - Field ID: 20116 and continuous - Field ID: 1239), batch, centre, and the first 40 genetic PCs. At this stage, trait residuals with an absolute skew *>* 2^[47]^ in the larger European ancestry group were removed to avoid measurement bias and the remaining trait values were standardised using a rank inverse normal transform on the residuals. This provided traits adjusted by key environmental factors, genetic PCs, and with relatively low skew, transformed to have standard normal distributions for all analyses.

#### Further Trait QC

To conduct our initial primary analyses, European ancestry UKB samples were split randomly into 50% base and 50% target data sets, for computing GWAS results and PRS (or “polygenic scores”), respectively. Then, for each trait, a GWAS was performed using the standardised trait values in the base data, applying PLINK-1.9^[48,49]^, while heritability and pairwise genetic correlations were estimated using LD score regression^[50]^. To ensure that our PRS analyses are well-powered, we removed traits with heritability *h*^2^ *<* 5%^[20]^, and to ensure that the traits were essentially polygenic, rather than oligogenic, traits were removed for which a single SNP (MAF > 1%) explained more than 2% of trait variance^[51]^. After these QC steps, 120 quantitative traits remained across all four categories.

To verify that the traits were meaningfully different from each other, we considered their pairwise genetic correlations. More than half (64 of the 120) of the traits had pairwise genetic correlation *r*_*g*_ *>* 95% with at least one other trait. Of these 64 traits, 22 had a single pairwise correlation above this threshold. For these 11 pairs of traits, the trait with higher heritability was selected for analysis. The remaining 42 genetically non-unique traits came from two categories: *Bone-densitometry of heel* and *Body composition by impedance*. Applying the same data maximisation algorithm used to select the maximum number of unrelated individuals^[45]^, we selected a further 7 traits (two bone-densitometry related and five body composition related) with pairwise *r*_*g*_ *<* 0.95. In addition to the 56 traits without high genetic pairwise correlations, and the 11 traits selected from correlated pairs, this resulted in a set of 74 traits for use in the primary analyses.

#### Initial PRS Analyses

The 74 traits remaining after QC included 37 in the *Biomarkers* category, 21 traits in *Physical Measures* category, and 16 that were in either the *Touchscreen Questionnaire* or *Cognitive Function* categories. In the primary unrelated European dataset, these traits had between of 50,109 and 167,857 (average 123,482) samples in each of the base and target datasets. For each (residualised) trait, PRSice-2^[52]^ was used to establish an optimal *P* -value threshold for computing PRS in the target data (2nd half of the data) using the *FastScore* option (10 thresholds tested). Each PRS and corresponding trait were then used in the POPout analyses, described below. While each PRS will be slightly overfit to the data due to training and testing within the same data, the use of only 10 *P* -value thresholds should minimise this overfitting, and overfitting of PRS to the trait should be independent, and in fact conservative, in relation to the results of the POPout analyses. This procedure optimised the sample size of the test data. None of the replication analyses (below) were subject to any overfitting since they all corresponded to out-of-sample data.

#### Replication Dataset QC

### European Ancestry Repeated Measures

After initial QC (above), there were 18,340 European ancestry individuals with measurements repeated at a follow-up data collection approximately an average of five years after baseline (timing differed for different individuals). For these individuals, both measurements were normalised alongside the other European ancestry UKB samples, as described above, but held out of the base dataset. Then the residualised trait values from their two visits were averaged (mean) to provide a target subset of individuals for which random measurement error, or transient effects e.g. infections, are less likely to produce trait values in the tail. Across the 74 traits, there were between 2,461 and 15,475 (mean: 10,699) unrelated individuals with repeated measures available for replication. PRS were calculated in this sample using the *P* -value threshold (and, thus, same SNPs) optimised for the relevant trait in the primary analyses, as well as the (same) effect size weights of the base GWAS.

### Multi-Ancestry Sample

After initial QC (above), there were 24,476 individuals of multiple non-European ancestries (see above) available for replication, which we refer to as the “Multi-Ancestry” replication sample. The trait values of these individuals were normalised alongside the European UKB individuals, as above, and PRS were calculated for the 74 traits of our primary analyses using PRSice-2^[52]^. PRS were calculated in this sample using the *P* -value threshold (and, thus, same SNPs) optimised for the relevant trait in the primary analyses, as well as the (same) effect size weights of the base GWAS. There were between 569 and 22,513 (average: 14,871) unrelated individuals for this multi-ancestry replication.

### All of Us Cohort

The All of Us Research Program (All of Us) is a diverse ancestry biobank from the United States, utilising electronic health record (EHR) data in a cohort of individuals with a wider age range than the UKB (*µ, σ*= 55.8,17.1 vs 56.5,8.1). Traits matching those analysed in our primary UKB analysis were identified through an initial database search and were then interrogated manually to determine if trait descriptions were sufficiently similar (see ***Supplementary Table 1***). All of Us provides ancestry categories, corresponding to definitions used within gnomAD^[53]^, the Human Genome Diversity Project^[54]^, and 1000 Genomes project^[55]^. 133,581 European ancestry individuals were extracted for analysis. Trait QC in All of Us was conducted to match that performed in the UKB as closely as possible, given available trait information. First of all, individuals on statins were removed from the *Blood count* and *Blood Biochemistry* subcategories, and then the median measurement value for every trait (EHR data includes multiple measures) was selected and the age corresponding to that measurement was extracted. For each trait, outliers greater than six standard deviations from the mean were removed and a linear regression model was applied with covariates for *age, age*^2^, *age*^3^, sex, and the first 40 genetic PCs. A rank inverse normal transformation was applied to the regression residuals to provide standardised trait values, as for the UKB.

Genetic QC in All of Us was conducted using standard filters, as used in the UKB. SNPs with MAF *<* 0.01, genotype missingness > 0.02, or Hardy-Weinberg equilibrium test *P <* 10^−8^ were excluded. The same greedy algorithm used to maximise sample retention of unrelated individuals as used in the UKB, was applied here. Using the higher-powered publicly available European ancestry GWAS summary statistics generated from the entire UKB^[35]^ to provide SNP weights, trait PRS were calculated^[52]^ using the same *P* -value threshold relevant to each trait and trained in the UKB European ancestry analysis (see above).

### Data Analysis: POPout Analyses

#### The POPout Test

Here we describe the PRS-On-Phenotype outlier test, or “POPout” test, which is designed to identify departures from polygenic expectations in the tails of quantitative traits from population genotype data. In our application, we seek to identify departures due to genetic factors, and, thus, we first perform a regression of traits on environmental risk factors (as described above) and use the residuals as input to the test. The POPout test first applies a regression of PRS on trait values, in which both the PRS and trait are standardised to have a standard deviation of 1. The assumption of the test is that, under uniform polygenicity, there is a linear relationship in prediction of PRS from trait values, but that under reduced polygenicity in the tails the fitted regression values systematically overestimate absolute PRS values in the lower and upper quantiles (i.e. observed PRS for extreme trait values will be closer to the population mean PRS than according to linear regression). The residuals from the regression are considered according to quantiles of the trait, and two-tailed t-tests are applied to test whether the residuals are significantly different from zero. Here we applied the test to individuals in the bottom and top 1% of the trait distribution to test for departures from polygenicity in the tails. The effect size in each tail is defined such that a positive effect is indicative of regression-to-the-mean in both the lower and upper tails, as follows:

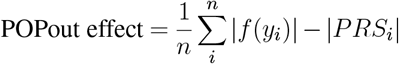

where, within the tested quantile, *f* (*y*) is the predicted PRS from the regression given trait value *y* and *n* is the number of observations.

#### POPout regression QC

An additional trait QC was applied to ensure the robustness of the POPout to its assumptions by ensuring that the POPout test did not produce significant results in the middle of the trait distribution. Specifically, for each trait, two-sided *t*-tests for zero mean were applied to the residuals assigned in each of the middle 80 centiles of the trait distribution. Traits were deemed to fail QC if the smallest *P* -value was significant at a Bonferroni-corrected level of 0.05 (*P* ≤ 0.05/80). Such traits were removed from our analysis.

#### Evaluating Impact of POPout Effects on PRS Prediction

To evaluate how observed POPout effects impact on the performance of polygenic score, or PRS, prediction, we emulate a scenario in which the quantitative trait underlies a binary outcome (e.g. as is the case for BMI underlying clinical obesity). We consider individuals above a specified trait threshold value to be cases and individuals below the threshold to be controls. We investigate thresholds at the 75% and the 99% points of the trait distribution. For each, we use the PRS to predict case/control status, and then generate a quantile plot, in which the odds ratio of the given quantile is estimated in relation to the central quantile via logistic regression (***Fig. 2c***). Additionally, for each trait with *PRS r*^2^, we simulated a quantitative phenotype with trait values given by *T* = *rPRS* + *E*, where *PRS* has a standard normal distribution and *E* represents all other contributions to trait variance and is drawn from a normal distribution with mean 0 and variance (1 − *r*^2^). This produces a simulated trait with a 𝒩 (0, 1) distribution and *PRS r*^2^. Using the same number of target samples used in the real data analysis for each trait, this simulation allows us to estimate the expected predictive performance of the PRS based on the corresponding *r*^2^ and assuming uniform polygenicity throughout the trait distribution.

### Replication of POPout Results

#### Bootstrap Study To Establish Power of Replication Analyses

In our primary analysis QC, we set a minimum sample size threshold of 100k in total (i.e. at least 50k target) samples. To explore whether our POPout results are expected to replicate with smaller sample sizes, we repeatedly sampled subsets of between 1k and 50k target samples and performed the POPout test in these subset or “bootstrap” samples. Using the POPout results in our primary analysis that were significant at FDR 5% as a ground truth, for each target sample of between 1k and 50k (incrementing by 5k), we performed 100 bootstraps across each of the 148 trait tails and counted the number that replication with (nominal P-value < 0.05). We also calculated the Pearson correlation between the bootstrap POPout effects and those of the full data at each bootstrap sample size. As a result of this investigation, we imposed a minimum replication target sample size of 5000 to provide sufficient power for replication (see ***Supplementary Fig. 1***).

#### Replication Across Datasets

We replicated our primary POPout analyses in three different independent cohorts: **(1)** UKB European ancestry individuals with repeat measures (baseline and follow-up at ∼5 years), **(2)** Non-European Multi-Ancestry UKB individuals, **(3)** the European ancestry samples in the US-based All of Us biobank. For each cohort, traits with at least 5000 samples that passed our POPout QC, were included and tested for matched discovery of POPout effects and correlation as described above.

### Sibling-based Analyses

#### Sibling-based Investigation of Departures from Polygenic Architecture

We have already developed sibling-based tests of departures from polygenic expectations in trait tails caused by *de novo* high-impact alleles or caused by rare alleles of large effect segregating in families (“Mendelian” effects)^[2]^. Here we summarise the theoretical framework underlying those tests, and then describe the novel test used here that combines the two tests into a single omnibus test to identify general departures from population expectations due to either cause.

The conditional distribution of an individual’s trait value given their sibling’s trait value, assuming the infinitesimal model for genetic effects (“polygenicity”) and normally distributed environmental factors, can be derived analytically^[2]^ as:

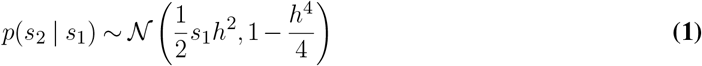

where *s*_1_ and *s*_2_ denote the index and conditional-sibling trait values, respectively, and *h*^2^ is the trait heritability. Rare large-effect alleles can produce deviations from this distribution, particularly when concentrated in specific parts of the distribution, which may occur in the trait tails if these effects are very large and abundant. Specifically, rare large-effect alleles segregating in families - *Mendelian effects* - will result in excess trait similarity (or concordance) between siblings on average compared to polygenicity. Conversely, a high-impact *de novo* mutation in one sibling will result in excess dissimilarity (discordance) between siblings.

Our sibling-based test for enrichment of *Mendelian* effects^[2]^ evaluates the proportion of sibling pairs in which both siblings reside in a defined quantile of the trait distribution, which can be formulated as a score test with the following z-statistic:

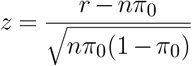

where *n* is the number of index siblings in the quantile *q* under test, *r* is the number of siblings pairs where both siblings have trait values in *q* and *π*_0_ is the expected proportion of sibling pairs in the quantile under polygenicity, which is given by:

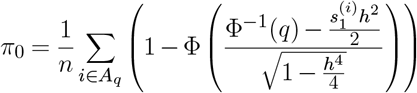

where *n* is the number of sibling pairs, *A*_*q*_ is the set of index siblings in trait quantile *q*, Φ is the normal cumulative density function and Φ^−1^ is its inverse.

Our sibling-based test for enrichment of *de novo* effects constructs a *z*-statistic for the distance between index siblings in the tail of the trait distribution and their conditional siblings, as follows:

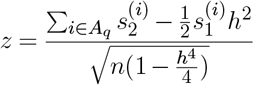

For index siblings in the lower and upper tails, the alternative hypotheses of *H*_1_ : *z >* 0 and *H*_1_ : *z <* 0 are tested as one-tailed *t*-tests, respectively. For further details, see Souaiaia et al^[2]^.

The trait heritability, *h*^2^, a required parameter in the *Mendelian* and *de novo* tests, is estimated from the sibling pairs by calculating the maximum likelihood of *h*^2^ under the null hypothesis of the infinitesimal model (see above):

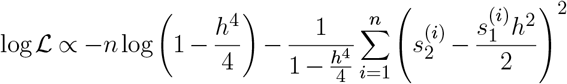

#### The Sibling STANDout Test

In order to test for general departures from polygenic architecture in the tails of quantitative traits, caused either by high-impact alleles that are either *de novo* or that are segregating in families (i.e. rare alleles), we combine the results of the two tests using Fisher’s method to give a *χ*^2^ statistic with 4 degrees of freedom:

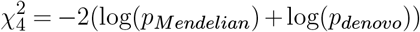

### Rare Variant and Selection Analyses

#### Associations between POPout effects and Rare Variants

Backman et al^[34]^ provides results from exome sequencing analysis performed in the UK Biobank full exome data across 3994 traits, including 65 of the 74 traits analysed in our target set. For each of these 65 traits, in each tail, the number of genes with at least one significant (P ≤ 2.18 × 10^−11^; significance threshold used in^[34]^) variant (deleterious, missense, predicted loss of function) or a genome-wide significant burden association is counted. Across the 130 trait tails, we calculate the Pearson correlation between the number of exome variants and the POPout effect size (***Fig. 4b***).

#### Relative Lifetime Reproductive Success

The number of offspring is recorded in the UKB for both females and males. Following the approach of Beauchamp^[56]^, samples were stratified into four birth cohorts (1934-42, 1943-48, 1949-55, 1956-65). Individual relative lifetime reproductive success, *Y*_*rLRS*_, was then calculated by dividing the number of offspring of each individual by the cohort mean. Following Sanjak et al^[30]^, the presence and direction of selective pressure acting on traits was inferred by fitting the following regression model:

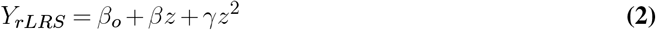

where *z* denotes the residualised and normalised trait values and *β* and *γ* are linear and quadratic regression parameters.

We employed Bayesian information criteria (BIC) to select the best fitting model for each trait, from which the selective pressure acting on each trait was inferred. If the selected model does not include a linear or quadratic parameter, then no selection is inferred. If the selected model includes only the linear term, then negative selection is inferred if *β <* 0, while positive selection is inferred if *β >* 0. If the selected model does not include a linear term but includes the quadratic term, with *γ <* 0, then stabilising selection is inferred, suggesting that the mean trait value in the population is optimal; if instead *γ >* 0, then this suggests that both tails of the distribution are subject to positive selection, which is known as “disruptive selection” and is expected to be rare in practice. When the BIC selected model includes both linear and quadratic terms, then the *β* parameter indicates the general direction of selection as positive or negative, with presence of the quadratic term suggesting that an optimal mean is being reached.

#### Associations between POPout Effects and LRS-inferred Selection

To test the association between POPout effects and the selection acting on corresponding traits as inferred by the rLRS model described above, we stratified traits into four categories of selection. The first two of the four categories reflect non-directional selection: **(1)** no selection (*β* = 0, *γ* = 0), in which POPout effects are not expected in either tail, **(2)** Non-directional stabilising selection (*β* = 0, *γ <* 0), in which we hypothesise POPout effects in both tails of equal magnitude. The third and fourth categories reflect directional selection: **(3)** positive selection (*β >* 0), which can be linear (*γ* = 0), disruptive ((*γ >* 0) or stabilising (*γ <* 0), in which we hypothesise greater POPout effects in the lower tail vs the upper tail, **(4)** negative selection (*β >* 0; linear, disruptive or stabilising as for positive), in which we hypothesise greater POPout effects in the upper tail vs the lower tail. For each category, a t-test is applied to test for the difference in mean POPout effects (excluding non-significant POPout effects) between the lower and upper tails across the traits.

#### Association between POPout Effects and Broad Measures of Fitness

Five broad measures of fitness were considered: **(1)** Male and **(2)** female fecundity (Field IDs: 2405 and 2734), **(3)** self-reported, non-cancer illnesses (Field ID: 2002), **(4)** total miscarriages and still births (Field IDs: 39290 and 3839), **(5)** paternal age (calculated as the difference between father’s age (Field ID: 2946) and participant age (Field ID: 21022). Associations between POPout effects and these measures were tested as follows. For each measure, the mean value was calculated separately in the upper and lower (1%) tail for every trait. Then, the association between between these tail measure means and corresponding POPout effect sizes was calculated as a Spearman’s rank correlation across the traits.

#### Population Genetics Forward-in-Time Simulation Study

Using SLiM v4.0 (1), we simulated a quantitative polygenic trait under a standard Wright-Fisher (WF) model of stabilising selection. First a population of N=10,000 diploid individuals of genome size 100kb was simulated to equilibrium (10N generations) with a constant mutation of 2.36 × 10^−8^ per base pair (bp) and recombination rate of 10^−8^ per bp under neutrality. Mutations were assigned effect sizes simulated from a gamma distribution with shape parameter 0.186 and mean 0.01026 and subsequently multiplied by -1 with probability 0.5 to give a symmetrical effect size distribution. This distribution has been used previously to simulate genetic effects in humans and reflects a prior belief in a small number of large effects and many small effects^[57]^. Individual trait values were determined by the sum of all genetic effects, thereby corresponding to full heritability (*h*^2^ = 1).

After neutral genetic diversity was achieved (10N generations), stabilising selection on the trait was introduced for 1N generations through a Gaussian fitness function

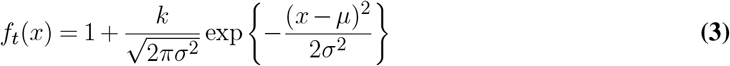

where *k* is a multiplier for strength of selection, *x* is the phenotype of an individual at generation *t, µ* and *σ*are the trait mean and standard deviation at the start of selection (i.e., *t* = 10*N*).

We ran this simulation for 100 independent replicates and reported data in each simulation by logging, at each generation, the phenotypic mean and standard deviation of the population, as well as statistics regarding the mutations present in the populations, including allele frequencies and effect sizes, and the mutational profile of each individual. To analyse the enrichment of rare variant architecture at different quantiles, the population was stratified across 100 centiles (each including 100 individuals) and the number of rare and common alleles carried by individuals in each centile was calculated under neutrality vs under selection. For comparison purposes, “weak selection” corresponded to simulations performed using a multiplier of *k* = 1 in equation 3, while “strong selection” corresponded to simulations with *k* = 100. “Large effects” were considered those in the top 10% of effect sizes genome-wide. The mean of the log-transformed odds ratio across all 100 independent simulation replicates was used as a measure of enrichment and a *t*-test was performed to test for significant differences from zero (see ***Fig. 4d***). To analyse the impact that low-frequency variants have on prediction and to mimic the calculation of common variant PRS in real data, a PRS was calculated using only variants with *MAF >* 1% and compared to the true trait value across the individuals in each centile. The mean difference (POPout effect) and 95% confidence intervals in each centile were calculated across the 100 independent simulation replicates (***Fig. 4e***). These simulations reflect a subset of those performed in a recent simulation study that we conducted^[58]^.

## Supplementary Material

**Supplementary Figure 1:**
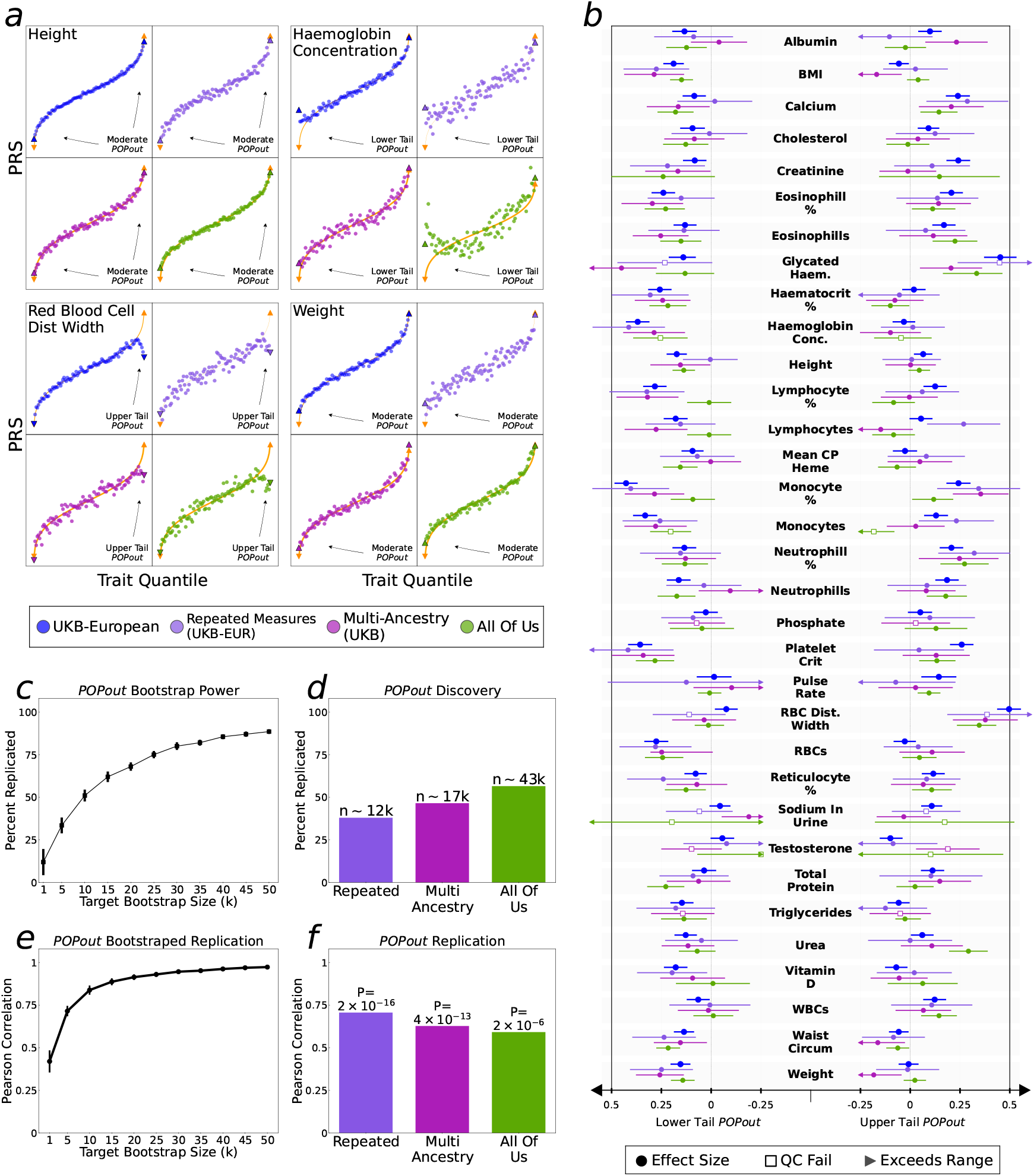
Extended Replication. **a**, PRS quantile (in 1% increments) plotted against mean trait value in quantile for four traits with different types of POPout effects in the primary analyses. **b**, POPout effect sizes in primary and replication cohorts across the 33 overlapping traits in the All of Us cohort. **c**,**d**, Considering trait tails with significant POPout results in our large European ancestry analysis (‘Discovery’) as true positives, the percent of tails replicated (*P* < 0.05) in 100 bootstrapped runs of different sizes (**c**) and in the replication datasets (**d**). **e**,**f**, Replication here is defined here as a significant Pearson correlation across effect sizes in our bootstrapped subsets (**e**) and in the replication datasets (**f**).

**Supplementary Table 1:**
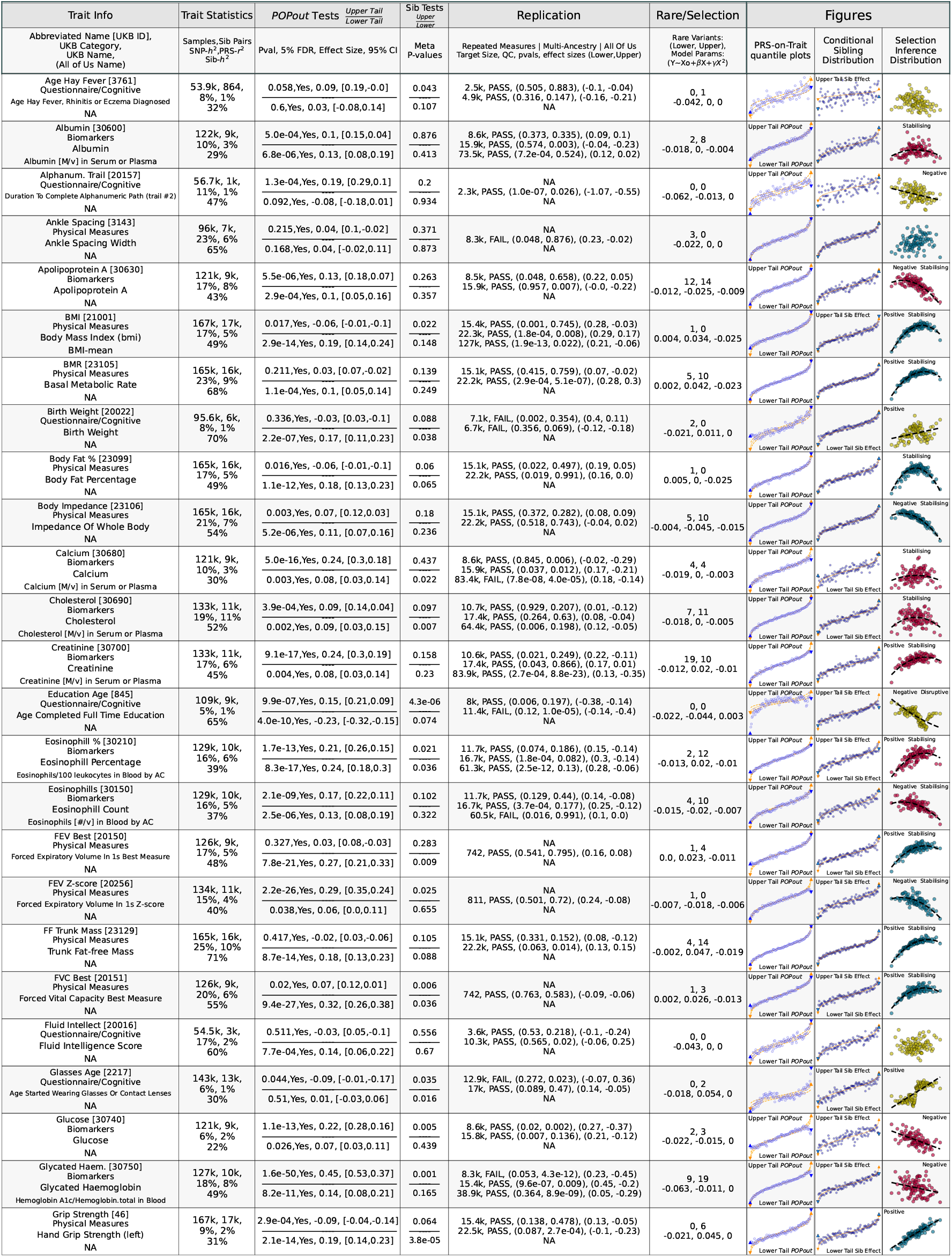
Traits 1-25.

**Supplementary Table 1:**
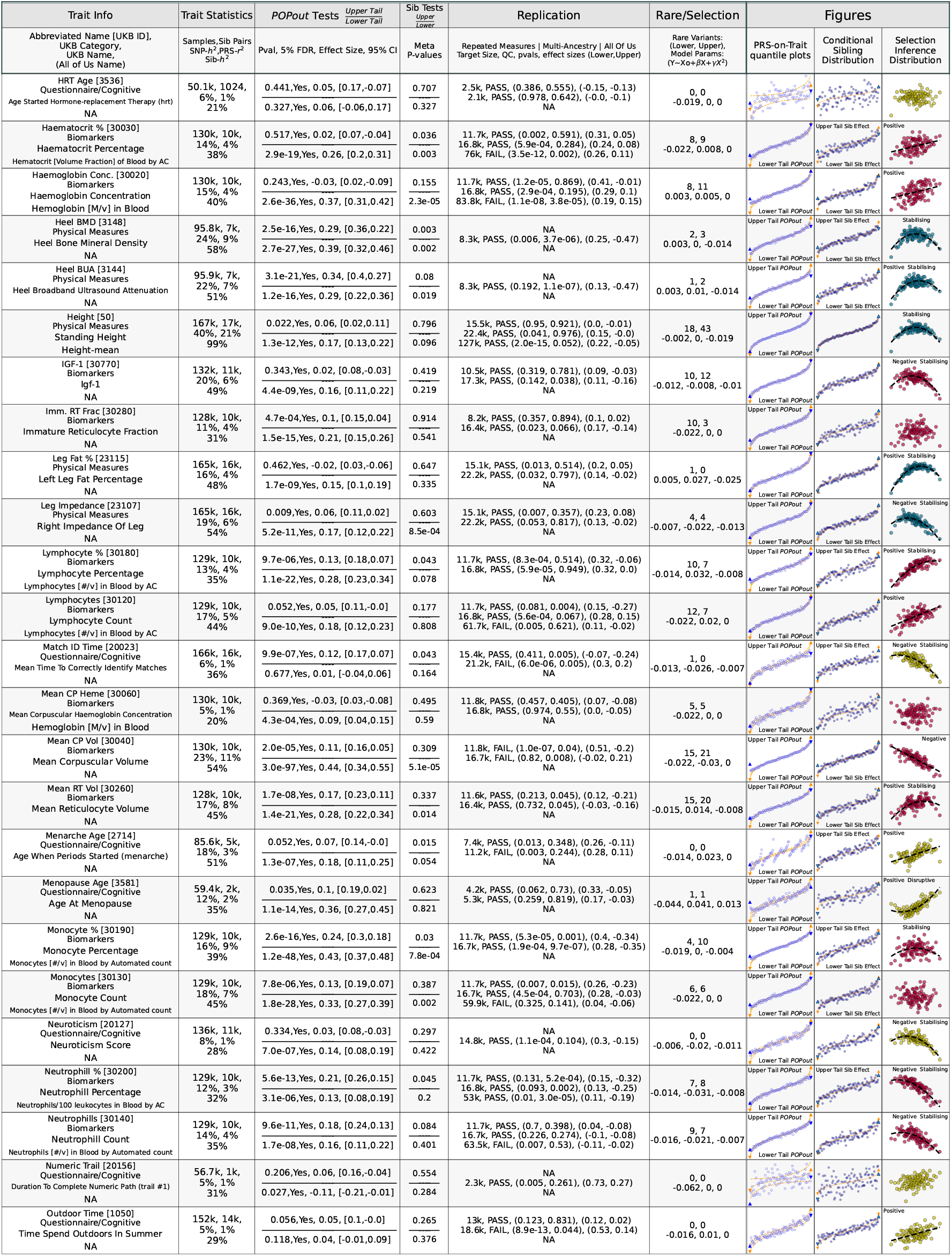
Traits 25-50.

**Supplementary Table 1:**
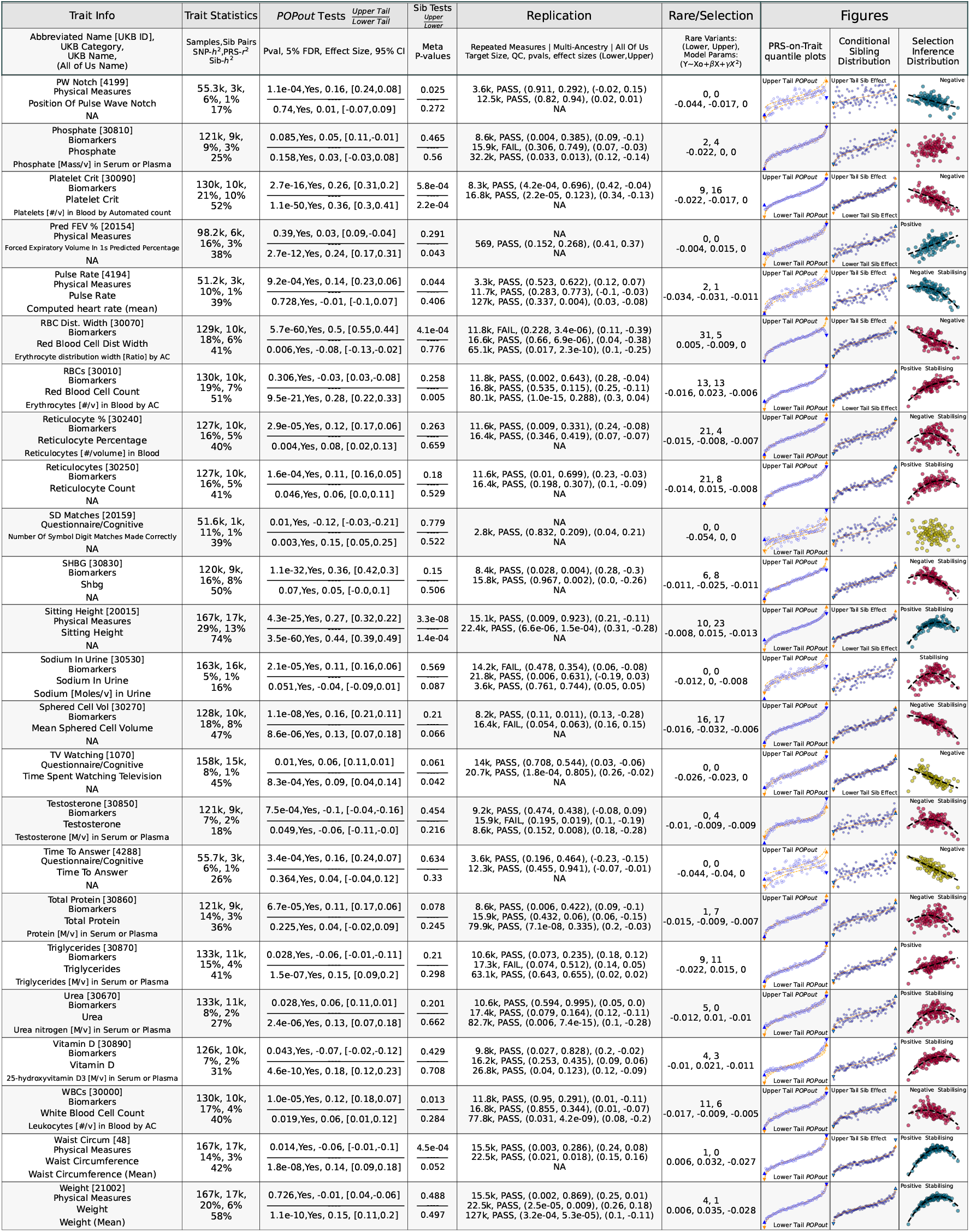
Traits 50-74.

## References

1. N. J. Timpson, C. M. T. Greenwood, N. Soranzo, D. J. Lawson, and J. B. Richards. Genetic architecture: the shape of the genetic contribution to human traits and disease. Nat. Rev. Genet., 19:110–124, 2018.

2. T. Souaiaia, H. M. Wu, C. Hoggart, and P. O’Reilly. Sibling similarity can reveal key insights into genetic architecture. eLife, 12, 2024.

3. Daniel J. Weiner, Ajay Nadig, Karthik A. Jagadeesh, Kushal K. Dey, Benjamin M. Neale, Elise B. Robinson, Konrad J. Karczewski, and Luke J. O’Connor. Polygenic architecture of rare coding variation across 394,783 exomes. Nature, 614(7948):492–499, 2023. doi: 10.1038/s41586-022-05684-z.

4. L.J. O’Connor et al. Extreme polygenicity of complex traits is explained by negative selection. Am. J. Hum. Genet., 105:456–476, 2019.

5. T. J. C. Polderman et al. Meta-analysis of the heritability of human traits based on fifty years of twin studies. Nat. Genet., 47:702–709, 2015.

6. A. Abdellaoui, L. Yengo, K. J. H. Verweij, and P. M. Visscher. 15 years of gwas discovery: Realizing the promise. Am. J. Hum. Genet., 110:179–194, 2023.

7. J. Flannick et al. Exome sequencing of 20,791 cases of type 2 diabetes and 24,440 controls. Nature, 570:71–76, 2019.

8. T. Singh et al. Rare coding variants in ten genes confer substantial risk for schizophrenia. Nature, 604:509–516, 2022.

9. L. Yengo et al. A saturated map of common genetic variants associated with human height. Nature, 610:704–712, 2022.

10. L. Corte, L. Liou, P. F. O’Reilly, and J. García-González. Trumpet plots: visualizing the relationship between allele frequency and effect size in genetic association studies. GigaByte, 2023:gigabyte89, 2023.

11. E. Koch et al. Genetic association data are broadly consistent with stabilizing selection shaping human common diseases and traits. Preprint, 2024. doi: 10.1101/2024.06.19.599789.

12. Yuval B. Simons, Kevin Bullaughey, Richard R. Hudson, and Guy Sella. A population genetic interpretation of gwas findings for human quantitative traits. PLOS Biology, 16(3):e2002985, 2018. doi: 10.1371/journal.pbio.2002985.

13. D. Golan, E. S. Lander, and S. Rosset. Measuring missing heritability: Inferring the contribution of common variants. Proc. Natl. Acad. Sci., 111:E5272–E5281, 2014.

14. P. Wainschtein et al. Assessing the contribution of rare variants to complex trait heritability from whole-genome sequence data. Nat. Genet., 54:263–273, 2022.

15. R. A. Fisher. The correlation between relatives on the supposition of mendelian inheritance. Earth Environ. Sci. Trans. R. Soc. Edinb., 52:399–433, 1919.

16. Naomi R. Wray, Cisca Wijmenga, Patrick F. Sullivan, Jian Yang, and Peter M. Visscher. Common disease is more complex than implied by the core gene omnigenic model. Cell, 173(7):1573–1580, 2018. doi: 10.1016/j.cell.2018.05.051.

17. N. H. Barton, A. M. Etheridge, and A. Véber. The infinitesimal model: Definition, derivation, and implications. Theor. Popul. Biol., 118:50–73, 2017.

18. J. Yang, S. H. Lee, M. E. Goddard, and P. M. Visscher. Gcta: A tool for genome-wide complex trait analysis. Am. J. Hum. Genet., 88:76–82, 2011.

19. B. K. Bulik-Sullivan et al. Ld score regression distinguishes confounding from polygenicity in genome-wide association studies. Nat. Genet., 47:291–295, 2015.

20. Shing Wan Choi, Timothy Shin-Heng Mak, and Paul F O’Reilly. Tutorial: a guide to performing polygenic risk score analyses. Nature protocols, 15 (9):2759–2772, 2020.

21. J. G. Kingsolver and D. W. Pfennig. Patterns and power of phenotypic selection in nature. BioScience, 57:561–572, 2007.

22. B. A. Ference, I. Graham, L. Tokgozoglu, and A. L. Catapano. Impact of lipids on cardiovascular health: Jacc health promotion series. J. Am. Coll. Cardiol., 72:1141–1156, 2018.

23. I. Kyrou, H. S. Randeva, C. Tsigos, G. Kaltsas, and M. O. Weickert. Clinical Problems Caused by Obesity. MDText.com, Inc., 2000.

24. M. van der Steen et al. Acan gene mutations in short children born sga and response to growth hormone treatment. J. Clin. Endocrinol. Metab., 102:1458–1467, 2017.

25. S. Saeed et al. Genetic variants in lep, lepr, and explain 30% of severe obesity in children from a consanguineous population. Obesity, 23:1687–1695, 2015.

26. Y. Miyachi, T. Miyazawa, and Y. Ogawa. Hnf1a mutations and beta cell dysfunction in diabetes. Int. J. Mol. Sci., 23:3222, 2022.

27. P. P. Fiziev et al. Rare penetrant mutations confer severe risk of common diseases. Science, 380:eabo1131, 2023.

28. C. Bycroft et al. The uk biobank resource with deep phenotyping and genomic data. Nature, 562:203–209, 2018.

29. A. G. Bick et al. Genomic data in the all of us research program. Nature, 627:340–346, 2024.

30. J. S. Sanjak, J. Sidorenko, M. R. Robinson, K. R. Thornton, and P. M. Visscher. Evidence of directional and stabilizing selection in contemporary humans. Proc. Natl. Acad. Sci., 115:151–156, 2018.

31. H. B. C. and P. W. Messer. Slim 3: Forward genetic simulation beyond the wright-fisher model. Mol. Biol. Evol., 36:632–637, 2019.

32. A. V. Khera et al. Genome-wide polygenic scores for common diseases identify individuals with risk equivalent to monogenic mutations. Nat. Genet., 50:1219–1224, 2018.

33. A. V. Khera et al. Polygenic prediction of weight and obesity trajectories from birth to adulthood. Cell, 177:587–596.e9, 2019.

34. Joshua D Backman, Alexander H Li, Anthony Marcketta, Dylan Sun, Joelle Mbatchou, Michael D Kessler, Christian Benner, Daren Liu, Adam E Locke, Suganthi Balasubramanian, et al. Exome sequencing and analysis of 454,787 uk biobank participants. Nature, 599:628–634, 2021.

35. Benjamin Neale. Neale lab data: http://www.nealelab.is/uk-biobank/, 2018.

36. Augustine Kong, Michael L Frigge, Gisli Masson, Soren Besenbacher, Patrick Sulem, Gisli Magnusson, Sigurjon A Gudjonsson, Asgeir Sigurdsson, Aslaug Jonasdottir, Adalbjorg Jonasdottir, et al. Rate of de novo mutations and the importance of father’s age to disease risk. Nature, 488(7412):471–475, 2012.

37. Seunggeun Lee, Mary J Emond, Michael J Bamshad, Kathleen C Barnes, Mark J Rieder, Deborah A Nickerson, David C Christiani, Mark M Wurfel, and Xihong Lin. Optimal unified approach for rare-variant association testing with application to small-sample case-control whole-exome sequencing studies. The American Journal of Human Genetics, 91(2):224–237, 2012.

38. Pardis C Sabeti, David E Reich, John M Higgins, Haninah ZP Levine, Daniel J Richter, Stephen F Schaffner, Stacey B Gabriel, Jill V Platko, Nick J Patterson, Gavin J McDonald, et al. Detecting recent positive selection in the human genome from haplotype structure. Nature, 419(6909):832–837, 2002.

39. Paul F O’Reilly, Ewan Birney, and David J Balding. Confounding between recombination and selection, and the ped/pop method for detecting selection. Genome research, 18(8):1304–1313, 2008.

40. Jonathan K Pritchard, Joseph K Pickrell, and Graham Coop. The genetics of human adaptation: hard sweeps, soft sweeps, and polygenic adaptation. Current biology, 20(4):R208–R215, 2010.

41. Ilse Höllinger, Pleuni S Pennings, and Joachim Hermisson. Polygenic adaptation: From sweeps to subtle frequency shifts. PLoS genetics, 15(3):e1008035, 2019.

42. Neda Barghi, Joachim Hermisson, and Christian Schlötterer. Polygenic adaptation: a unifying framework to understand positive selection. Nature Reviews Genetics, 21(12):769–781, 2020.

43. Cathie Sudlow, John Gallacher, Naomi Allen, Valerie Beral, Paul Burton, John Danesh, Paul Downey, Paul Elliott, Jane Green, Martin Landray, et al. Uk biobank: an open access resource for identifying the causes of a wide range of complex diseases of middle and old age. PLoS medicine, 12(3):e1001779, 2015.

44. Information for Researchers. Ukb genotyping and quality control (v.1.2): https://biobank.ctsu.ox.ac.uk/crystal/crystal/docs/genotyping_qc.pdf, 2015.

45. Shing Wan Choi, Timothy Shin Heng Mak, Clive J Hoggart, and Paul F O’Reilly. Erasor: a software tool to eliminate inflation caused by sample overlap in polygenic score analyses. GigaScience, 12:giad043, 2023.

46. Rosa Cheesman, Jonathan Coleman, Christopher Rayner, KL Purves, Genevieve Morneau-Vaillancourt, Kylie Glanville, Shing W Choi, Gerome Breen, and Thalia C Eley. Familial influences on neuroticism and education in the uk biobank. Behavior genetics, 50(2):84–93, 2020.

47. Paul Mallery and Darren George. SPSS for windows step by step. Allyn & Bacon, Inc., 2000.

48. Shaun Purcell, Benjamin Neale, Kathe Todd-Brown, Lori Thomas, Manuel AR Ferreira, David Bender, Julian Maller, Pamela Sklar, Paul IW De Bakker, Mark J Daly, et al. Plink: a tool set for whole-genome association and population-based linkage analyses. The American journal of human genetics, 81(3):559–575, 2007.

49. Christopher C Chang, Carson C Chow, Laurent CAM Tellier, Shashaank Vattikuti, Shaun M Purcell, and James J Lee. Second-generation plink: rising to the challenge of larger and richer datasets. Gigascience, 4(1):s13742–015, 2015.

50. Guiyan Ni, Gerhard Moser, Stephan Ripke, Benjamin M Neale, Aiden Corvin, James TR Walters, Kai-How Farh, Peter A Holmans, Phil Lee, Brendan Bulik-Sullivan, et al. Estimation of genetic correlation via linkage disequilibrium score regression and genomic restricted maximum likelihood. The American Journal of Human Genetics, 102(6):1185–1194, 2018.

51. Arthur Korte, Bjarni J Vilhjálmsson, Vincent Segura, Alexander Platt, Quan Long, and Magnus Nordborg. A mixed-model approach for genome-wide association studies of correlated traits in structured populations. Nature genetics, 44(9):1066–1071, 2012.

52. Shing Wan Choi and Paul F O’Reilly. Prsice-2: Polygenic risk score software for biobank-scale data. Gigascience, 8(7):giz082, 2019.

53. Konrad J Karczewski, Laurent C Francioli, Grace Tiao, Beryl B Cummings, Jessica Alföldi, Qingbo Wang, Ryan L Collins, Kristen M Laricchia, Andrea Ganna, Daniel P Birnbaum, et al. The mutational constraint spectrum quantified from variation in 141,456 humans. Nature, 581(7809):434–443, 2020.

54. L Luca Cavalli-Sforza. The human genome diversity project: past, present and future. Nature Reviews Genetics, 6(4):333–340, 2005.

55. 1000 Genomes Project Consortium. A map of human genome variation from population-scale sequencing. Nature, 467(7319):1061–1073, 2010.

56. Jonathan P Beauchamp. Genetic evidence for natural selection in humans in the contemporary united states. Proceedings of the National Academy of Sciences, 113(28):7774–7779, 2016.

57. Xin He, Stephan J Sanders, Li Liu, Silvia De Rubeis, Elaine T Lim, James S Sutcliffe, Gerard D Schellenberg, Richard A Gibbs, Mark J Daly, Joseph D Buxbaum, et al. Integrated model of de novo and inherited genetic variants yields greater power to identify risk genes. PLoS genetics, 9(8):e1003671, 2013.

58. A. P. S. Ori, C. Giner-Delgado, C. J. Hoggart, and P.F. O’Reilly. Stabilising selection enriches the tails of complex traits with rare alleles of large effect. Preprint, 2024. doi: 10.1101/2024.09.12.612687.

